## Supplementary Materials for "Nucleotide context models outperform protein language models for predicting antibody affinity maturation"

| Data set | Species | PCPs | Clonal families | Median PCPs/CF | Median subs./PCP | Comments |
|---|---|---|---|---|---|---|
| Tang et al. IgH [55] | Human | 438,302 | 130,621 | 2 | 2 | In OAS (AbLang training set) |
| Jaffe et al. IgH [31] | Human | 176,656 | 49,430 | 2 | 4 | In OAS (AbLang training set) and Thrifty-prod training set |
| Rodriguez et al. IgH [30] | Human | 18,639 | 6,837 | 2 | 4 | Not in OAS or any training set |
| Ford et al. IgH [29] | Human | 2,407 | 1,182 | 2 | 3 | Not in OAS or any training set |
| Replay IgH [32] | Mouse | 5,378 | 129 | 44 | 1 | Not in OAS or any training set |
| Replay Ig$\kappa$ [32] | Mouse | 5,176 | 129 | 40 | 1 | Not in OAS or any training set |

**Table S1. Summary of data sets used.** All human data results in the main text are based on the Tang et al. IgH data set.

tab:pcp˙stats

| Model | Overlap | R-prec. | Sub. Acc. | CSP Perp. |
|---|---|---|---|---|
| S5F | 0.782 | 0.118 | 0.263 | 8.20 |
| S5F + ESM-1v | 0.617 | 0.112 | 0.328 | 9.12 |
| S5F + BLOSUM62 | 0.795 | 0.122 | 0.300 | 7.84 |
| Thrifty-SHM | 0.823 | 0.145 | 0.326 | 6.87 |
| Thrifty-SHM + ESM-1v | 0.624 | 0.116 | 0.334 | 8.63 |
| Thrifty-SHM + BLOSUM62 | 0.835 | 0.149 | 0.350 | 6.67 |
| Thrifty-prod | **0.900** | **0.163** | **0.370** | **6.36** |
| ESM-1v | 0.739 | 0.091 | 0.252 | 13.48 |
| AbLang2 | 0.780 | 0.095 | 0.353 | 8.20 |

**Table S2. Model performance on Rodriguez et al. IgH data set.** The best model for each metric is indicated in **bold**. The Rodriguez et al. data set is not part of any model training data. tab:rodriguez˜results

| Model | Overlap | R-prec. | Sub. Acc. | CSP Perp. |
|---|---|---|---|---|
| S5F | 0.779 | 0.106 | 0.261 | 8.33 |
| S5F + ESM-1v | 0.613 | 0.096 | 0.324 | 9.61 |
| S5F + BLOSUM62 | 0.790 | 0.107 | 0.300 | 8.01 |
| Thrifty-SHM | 0.824 | 0.128 | 0.324 | 7.00 |
| Thrifty-SHM + ESM-1v | 0.620 | 0.102 | 0.328 | 8.99 |
| Thrifty-SHM + BLOSUM62 | 0.834 | 0.132 | 0.345 | 6.80 |
| Thrifty-prod | **0.900** | **0.141** | **0.365** | **6.52** |
| ESM-1v | 0.733 | 0.082 | 0.249 | 13.72 |
| AbLang2 | 0.787 | 0.090 | 0.332 | 8.61 |

**Table S3. Model performance on Ford et al. IgH data set.** The best model for each metric is indicated in **bold**. The Ford et al. data set is not part of any model training data. tab:ford˜results

| Model | Overlap | R-prec. | Sub. Acc. | CSP Perp. |
|---|---|---|---|---|
| S5F | 0.783 | 0.096 | 0.262 | 8.19 |
| S5F + ESM-1v | 0.616 | 0.100 | 0.326 | 9.11 |
| S5F + BLOSUM62 | 0.786 | 0.098 | 0.299 | 7.92 |
| Thrifty-SHM | 0.834 | 0.127 | 0.328 | 6.86 |
| Thrifty-SHM + ESM-1v | 0.627 | 0.103 | 0.331 | 8.65 |
| Thrifty-SHM + BLOSUM62 | 0.836 | 0.129 | 0.344 | 6.73 |
| Thrifty-prod | **0.923** | **0.139** | **0.363** | **6.46** |
| ESM-1v | 0.744 | 0.082 | 0.255 | 12.61 |
| AbLang2 | 0.774 | 0.097 | 0.356 | 7.61 |

**Table S4. Model performance on the Tang et al. IgH data set.** The best model for each metric is indicated in **bold**. The Tang et al. data set is part of the AbLang2 training data. tab:tang˜results

| Model | Overlap | R-prec. | Sub. Acc. | CSP Perp. |
|---|---|---|---|---|
| S5F | 0.771 | 0.111 | 0.277 | 7.38 |
| S5F + ESM-1v | 0.595 | 0.099 | 0.341 | 8.10 |
| S5F + BLOSUM62 | 0.780 | 0.113 | 0.319 | 7.01 |
| Thrifty-SHM | 0.818 | 0.147 | 0.347 | 6.22 |
| Thrifty-SHM + ESM-1v | 0.603 | 0.107 | 0.345 | 7.73 |
| Thrifty-SHM + BLOSUM62 | 0.823 | 0.150 | 0.369 | 6.00 |
| Thrifty-prod | **0.907** | **0.164** | 0.393 | **5.66** |
| ESM-1v | 0.710 | 0.073 | 0.258 | 12.43 |
| AbLang2 | 0.754 | 0.086 | **0.41** | 6.23 |

**Table S5. Model performance on Jaffe et al. IgH data set.** The best model for each metric is indicated in **bold**. The Jaffe et al. data set is the training data for Thrifty-prod and part of the training data for AbLang2.tab:jaffe˙results

| Model | Modeling perspective | Biological process | EPAM framework | Species evaluated in EPAM |
|---|---|---|---|---|
| Thrifty-SHM [16] | ME | SHM | NT | Human |
| Thrifty-prod [16] | ME | Implicit SHM-selection | NT | Human |
| S5F [13] | ME | SHM | NT | Human |
| S5F + ESM-1v | ME+LM | Explicit SHM-selection | NT-AA | Human |
| S5F + BLOSUM62 [61] | ME | Explicit SHM-selection | NT-AA | Human |
| Thrifty-SHM + ESM-1v | ME+LM | Explicit SHM-selection | NT-AA | Human |
| Thrifty-SHM + BLOSUM62 | ME | Explicit SHM-selection | NT-AA | Human |
| ESM-1v (model 1) [35] | LM | Selection | AA | Human, mouse |
| AbLang [21, 28] | LM | Implicit SHM-selection | AA | Human, mouse |
| ReplaySHM [32] | ME | SHM | NT | Mouse |
| ReplaySHM + DMS | ME | Explicit SHM-selection | NT-AA | Mouse |
| ReplaySHM + BLOSUM62 | ME | Explicit SHM-selection | NT-AA | Mouse |
| ReplaySHM + ESM-1v | ME+LM | Explicit SHM-selection | NT-AA | Mouse |

**Table S6. Classifications for all models considered.** Models are characterized by the modeling perspective considered: molecular evolution (ME) or language modeling (LM). The molecular evolution perspective includes mutation models (SHM) and models that combine mutation and selection (explicit SHM-selection), while language models either characterize the joint substitution process (implicit SHM-selection) or selection alone. EPAM frameworks are classified by the input data type, considering nucleotide (NT) and/or amino acid (AA) sequences. Models are evaluated on human and mouse data sets in EPAM where applicable.

tab:model˙classifications

| | 23 | 41 | 43 | 98 | 102 | 104 | 118 |
|---|---|---|---|---|---|---|---|
| Observed | 0.14 | 0.15 | 0.13 | 0.19 | 0.25 | 0 | 0 |
| S5F | 0.77 | 0.46 | 0.33 | 0.60 | 0.98 | 0.72 | 0.43 |
| Thrifty-SHM | 0.54 | 0.51 | 0.39 | 0.49 | 0.85 | 0.61 | 0.43 |
| ESM-1v | 0.11 | 0.11 | 0.15 | 0.11 | 0.14 | 0.11 | 0.16 |
| Thrifty-SHM + ESM-1v | 0.09 | 0.09 | 0.10 | 0.10 | 0.11 | 0.09 | 0.10 |
| AbLang2 | 0.12 | 0.27 | 0.25 | 0.30 | 0.46 | 0.12 | 0.39 |
| Thrifty-prod | 0.27 | 0.27 | 0.22 | 0.23 | 0.49 | 0.27 | 0.23 |

**Table S7. Quantile of the observed or expected number of substitutions at known conserved sites in the Tang et al. IgH data set.**

tab:conserved˙sites˙quantile

| Model | Overlap | R-prec. | Sub. Acc. | CSP Perp. |
|---|---|---|---|---|
| ReplaySHM | 0.790 | 0.143 | 0.416 | 5.07 |
| ReplaySHM + BLOSUM62 | 0.777 | 0.129 | 0.420 | **5.01** |
| ReplaySHM + DMS | **0.820** | **0.146** | **0.430** | 5.10 |
| ReplaySHM + ESM-1v | 0.535 | 0.034 | 0.374 | 7.69 |
| ESM-1v | 0.530 | 0.011 | 0.212 | 15.9 |
| AbLang2 | 0.623 | 0.023 | 0.385 | 5.88 |

**Table S8. Model performance on Replay IgH data set.** The best model for each metric is indicated in **bold**.

tab:gcreplay˙igh˙results

| Model | Overlap | R-prec. | Sub. Acc. | CSP Perp. |
|---|---|---|---|---|
| ReplaySHM | 0.717 | 0.090 | 0.386 | 4.91 |
| ReplaySHM + BLOSUM62 | 0.710 | 0.091 | 0.419 | 4.79 |
| ReplaySHM + DMS | **0.786** | **0.125** | **0.423** | **4.77** |
| ReplaySHM + ESM-1v | 0.550 | 0.039 | 0.377 | 8.70 |
| ESM-1v | 0.547 | 0.017 | 0.182 | 21.24 |
| AbLang2 | 0.646 | 0.022 | 0.367 | 7.30 |

**Table S9. Model performance on Replay Ig$\kappa$ data set.** The best model for each metric is indicated in **bold**.

tab:gcreplay˙igk˙results

| DMS data | chain | Overlap | R-prec. | Sub. Acc. | CSP Perp. |
|---|---|---|---|---|---|
| bind | IgH | 0.820 | 0.146 | 0.430 | 5.10 |
| expr | IgH | 0.797 | 0.143 | 0.420 | 5.06 |
| bind+expr | IgH | 0.821 | 0.146 | 0.430 | 5.09 |
| bind | Ig$\kappa$ | 0.786 | 0.125 | 0.423 | 4.77 |
| expr | Ig$\kappa$ | 0.725 | 0.090 | 0.386 | 4.88 |
| bind+expr | Ig$\kappa$ | 0.789 | 0.125 | 0.423 | 4.77 |

**Table S10. Incorporating expression measurements for ReplaySHM+DMS.** Performance comparisons for different sources of DMS measurements to compute selection factors for the ReplaySHM + DMS model. For binding measurements (bind), as reported in the main results, selection factors are the ratio of dissociation contants, $R = K_D^{\text{parent}}/K_D^{\text{child}}$. For expression measurements (expr), the selection factors are the ratio of expression values, $R = E^{\text{child}}/E^{\text{parent}}$. For combining binding and expression (bind+expr), the selection factors are computed as, $R = (K_D^{\text{parent}}/K_D^{\text{child}})\,(E^{\text{child}}/E^{\text{parent}})$. tab:gcreplay˙expr

| | IgH | | | Ig$\kappa$ | | |
|---|---|---|---|---|---|---|
| | naive | leaf | all | naive | leaf | all |
| No. of PCPs | 391 | 4096 | 5378 | 424 | 3772 | 5176 |
| No. of amino acid substitutions | 595 | 6772 | 8499 | 702 | 6094 | 8107 |
| overlap: ReplaySHM | 0.654 | 0.822 | 0.790 | 0.545 | 0.751 | 0.717 |
| overlap: ReplaySHM + DMS | 0.649 | 0.847 | 0.820 | 0.580 | 0.820 | 0.786 |
| overlap: relative change (%) | -0.8 | +3.0 | +3.7 | +6.6 | +9.2 | +9.6 |
| R-prec: ReplaySHM | 0.161 | 0.128 | 0.143 | 0.045 | 0.088 | 0.090 |
| R-prec: ReplaySHM + DMS | 0.158 | 0.131 | 0.146 | 0.103 | 0.118 | 0.125 |
| R-prec: relative change (%) | -1.8 | +2.4 | +2.4 | +126 | +33 | +38 |
| sub. acc.: ReplaySHM | 0.316 | 0.411 | 0.416 | 0.281 | 0.406 | 0.386 |
| sub. acc.: ReplaySHM + DMS | 0.365 | 0.422 | 0.430 | 0.400 | 0.423 | 0.423 |
| sub. acc.: relative change (%) | +15 | +2.7 | +3.4 | +43 | +4.2 | +9.5 |
| CSP perp.: ReplaySHM | 6.324 | 5.035 | 5.070 | 5.543 | 4.859 | 4.907 |
| CSP perp.: ReplaySHM + DMS | 5.116 | 5.269 | 5.096 | 4.673 | 4.951 | 4.772 |
| CSP perp.: change | -1.207 | +0.234 | +0.026 | -0.870 | +0.092 | -0.135 |

**Table S11. Performance comparisons of ReplaySHM versus ReplaySHM + DMS on naive-parent PCPs and leaf-child PCPs.** The data set is split into PCPs with parent sequence being the naive sequence (the "naive" columns) and PCPs with child sequence being a leaf node (the "leaf" columns). Most of the data are in leaf-child PCPs. Performance metrics for the ReplaySHM model and the ReplaySHM + DMS model are reported, with change in performance compared to ReplaySHM. The "relative change" is the difference in the metric divided by the ReplaySHM value, reported as a percentage. For CSP perplexity, the absolute difference is reported; a lower CSP perplexity score is better and a negative difference indicates an improvement.

tab:gcreplay˙naive˙leaf˙summary

| **Model** | **IgH aff.** | **Ig$\kappa$ aff.** | **IgH expr.** | **Ig$\kappa$ expr.** |
|---|---|---|---|---|
| ESM-1v 1 | 0.0317 | 0.2636 | 0.3701 | 0.5538 |
| ESM-1v 2 | 0.0466 | 0.2677 | 0.4016 | 0.5739 |
| ESM-1v 3 | 0.0438 | 0.2414 | 0.3890 | 0.5284 |
| ESM-1v 4 | 0.0527 | 0.2427 | 0.4163 | 0.5688 |
| ESM-1v 5 | 0.0421 | 0.2551 | 0.3788 | 0.5699 |
| ESM-1v ensemble | 0.0477 | 0.2658 | 0.4076 | 0.5793 |

**Table S12. DMS and ESM-1v correlations for Replay IgH and Ig$\kappa$ naive sequences.** The ESM-1v values used here are the log transformed probability ratios (site-specific probabilities relative to the wildtype amino acid). Correlation values represent the Spearman rank correlation coefficient between DMS affinity or expression measures and ESM-1v predictions. We observed a stronger correlation between ESM-1v and DMS expression values relative to affinity. Similar trends were found in large benchmarks of DMS measurements and ESM models [40]. tab:model˙affinities

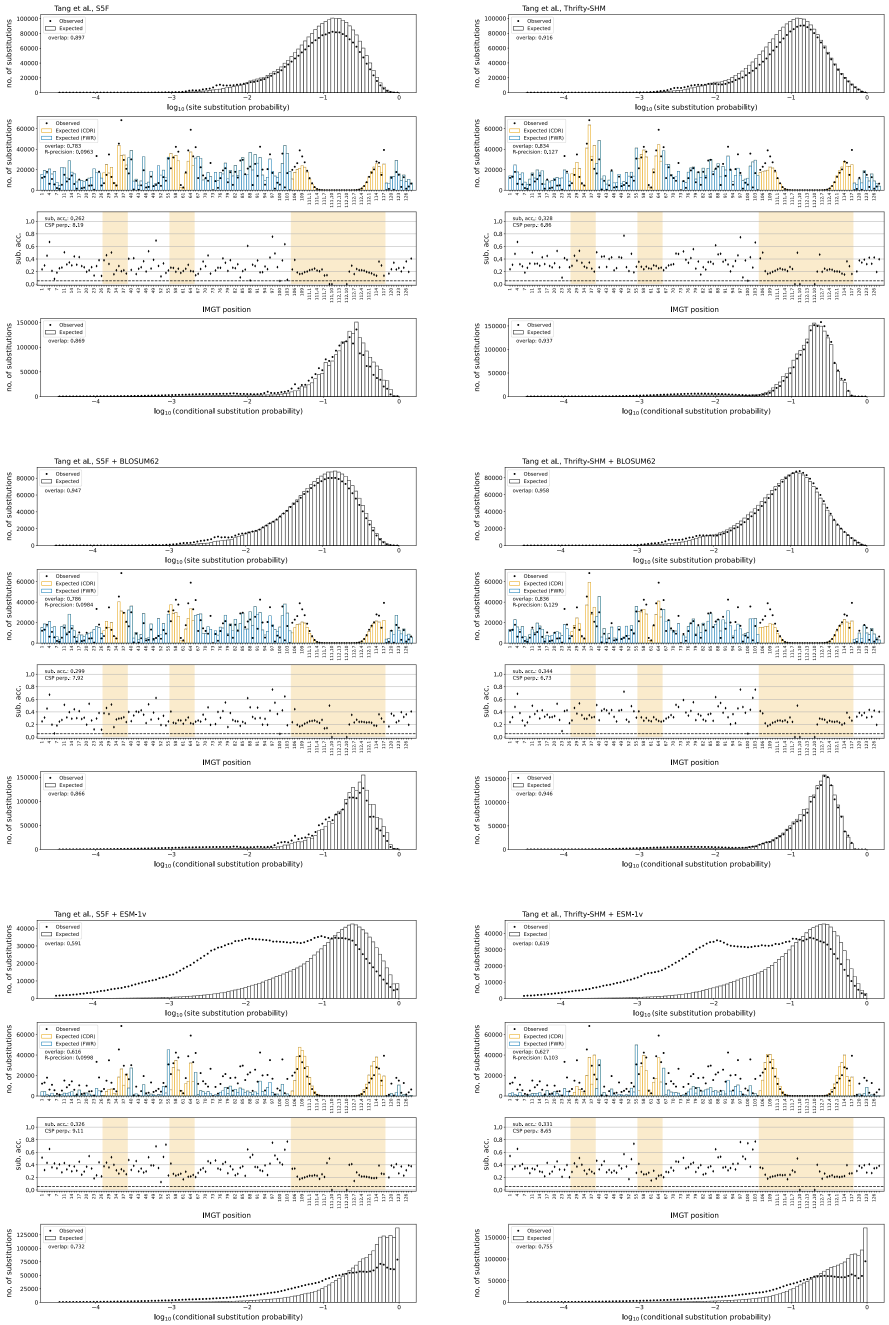


**Fig S1. NT and NT-AA framework models on the Tang et al. data set.** For each model, there are four panels. The top panel shows the observed and expected number of substitutions where the sites are binned according to their predicted SSP. The second panel shows the observed and expected number of substitutions over site position. The third panel shows the per-site substitution accuracy. The bottom panel shows the observed and expected number of substitutions at sites of substitution, and the sites are binned according to their predicted CSP. The hatch-filled bands are the standard deviations on the expectations.

fig:tang˙all˙1

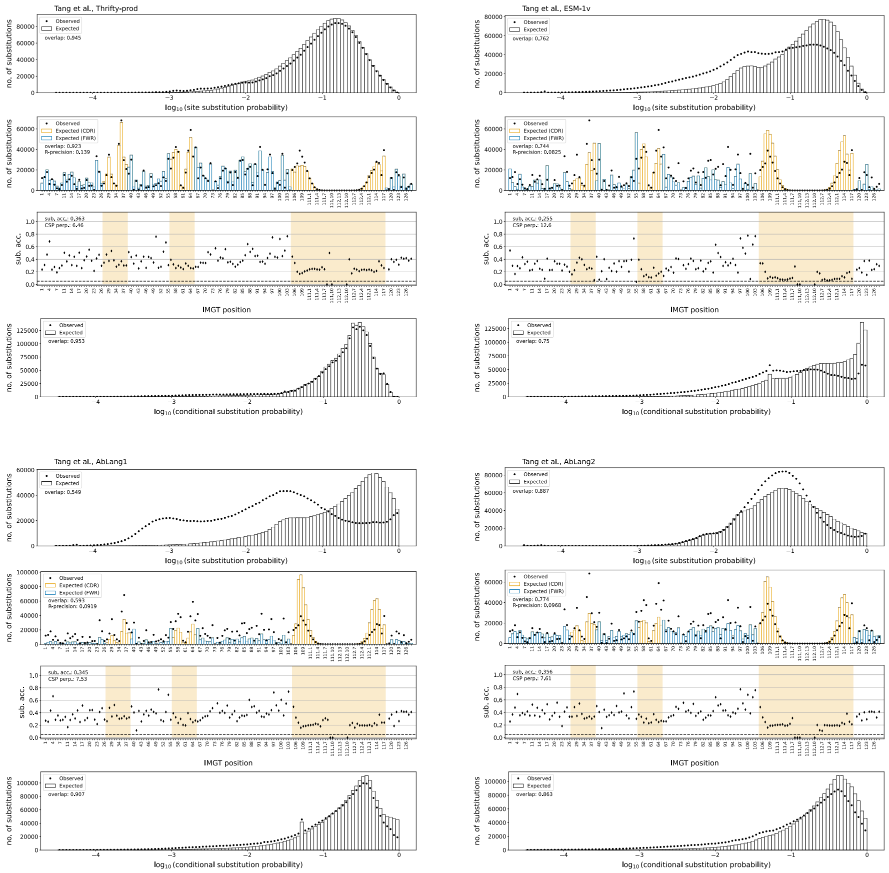


**Fig S2. AA framework models on the Tang et al. data set.** For each model, there are four panels. The top panel shows the observed and expected number of substitutions where the sites are binned according to their predicted SSP. The second panel shows the observed and expected number of substitutions over site position. The third panel shows the per-site substitution accuracy. The bottom panel shows the observed and expected number of substitutions at sites of substitution, and the sites are binned according to their predicted CSP. The hatch-filled bands are the standard deviations on the expectations.

fig:tang˙all˙2

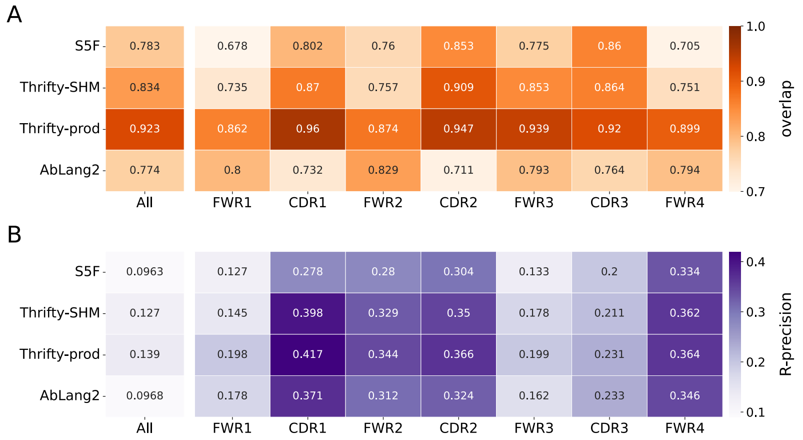


**Fig S3. Per-region overlap and R-precision on the Tang et al. data set.** (A) Site substitution overlap. (B) R-precision.

fig:human˙overlap˙and˙rprec

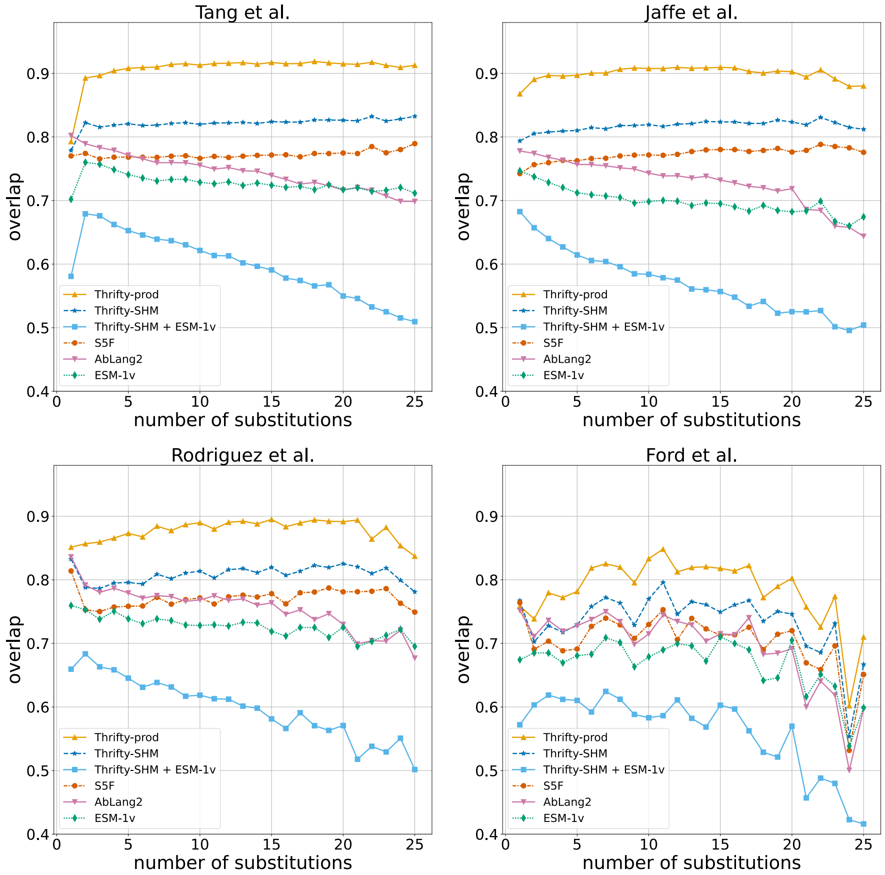


**Fig S4. Overlap stratified by number of substitutions per PCP evaluated on the human data sets.**

fig:human˙overlap˙vs˙nmuts

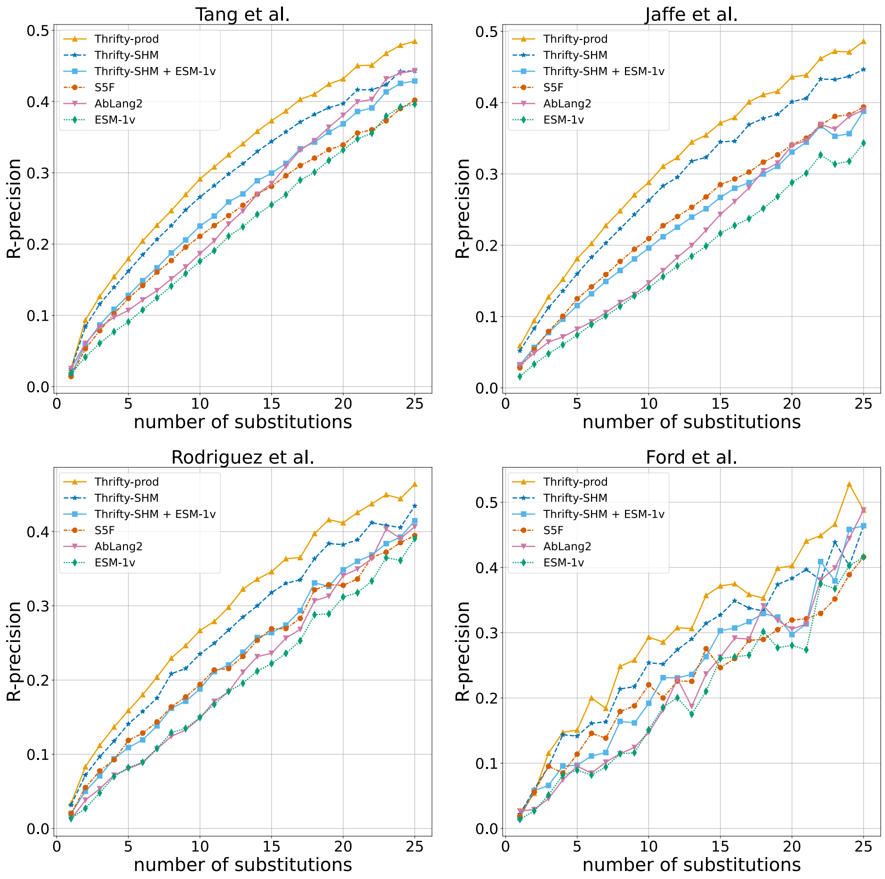


**Fig S5. R-precision stratified by number of substitutions per PCP evaluated on the human data sets.**

fig:human`rprec`vs`nmuts

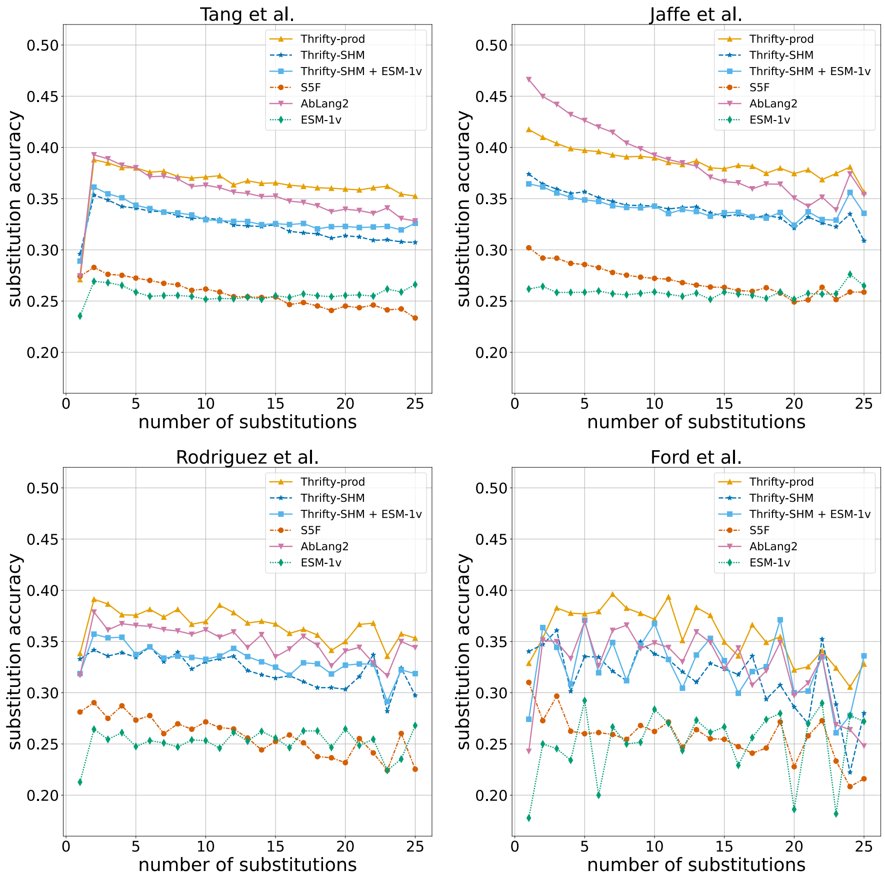


**Fig S6. Substitution accuracy stratified by number of substitutions per PCP evaluated on the human data sets.** fig:human`subacc`vs`nmuts

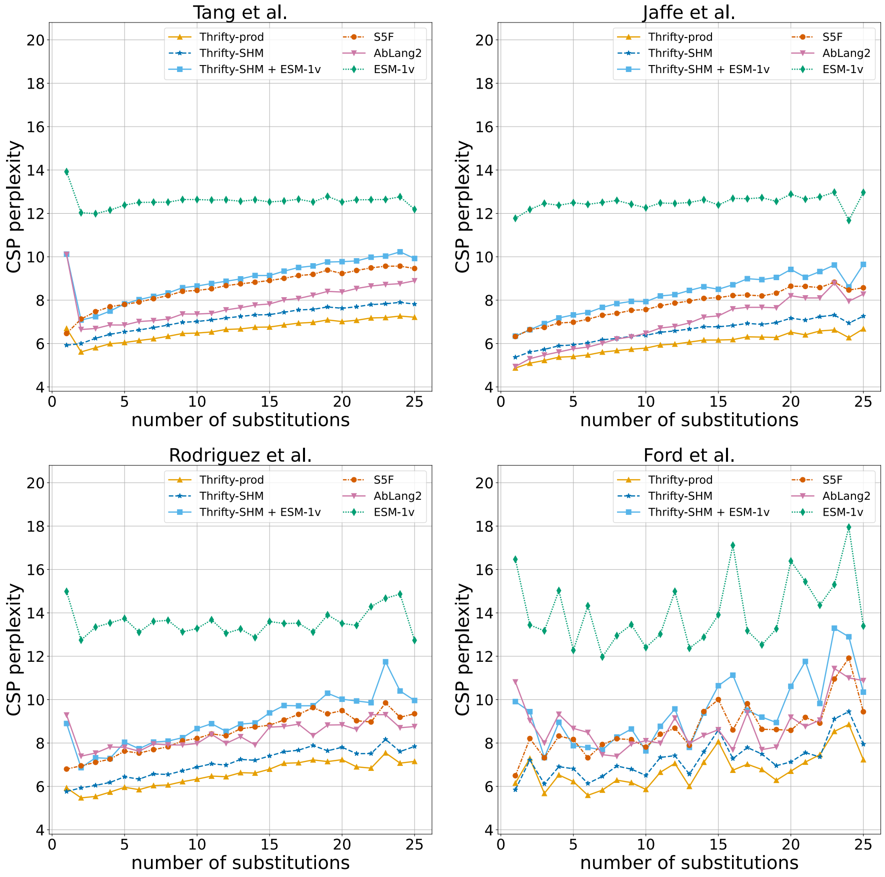


**Fig S7. CSP perplexity stratified by number of substitutions per PCP evaluated on the human data sets.**

fig:human`cspperp`vs`nmuts

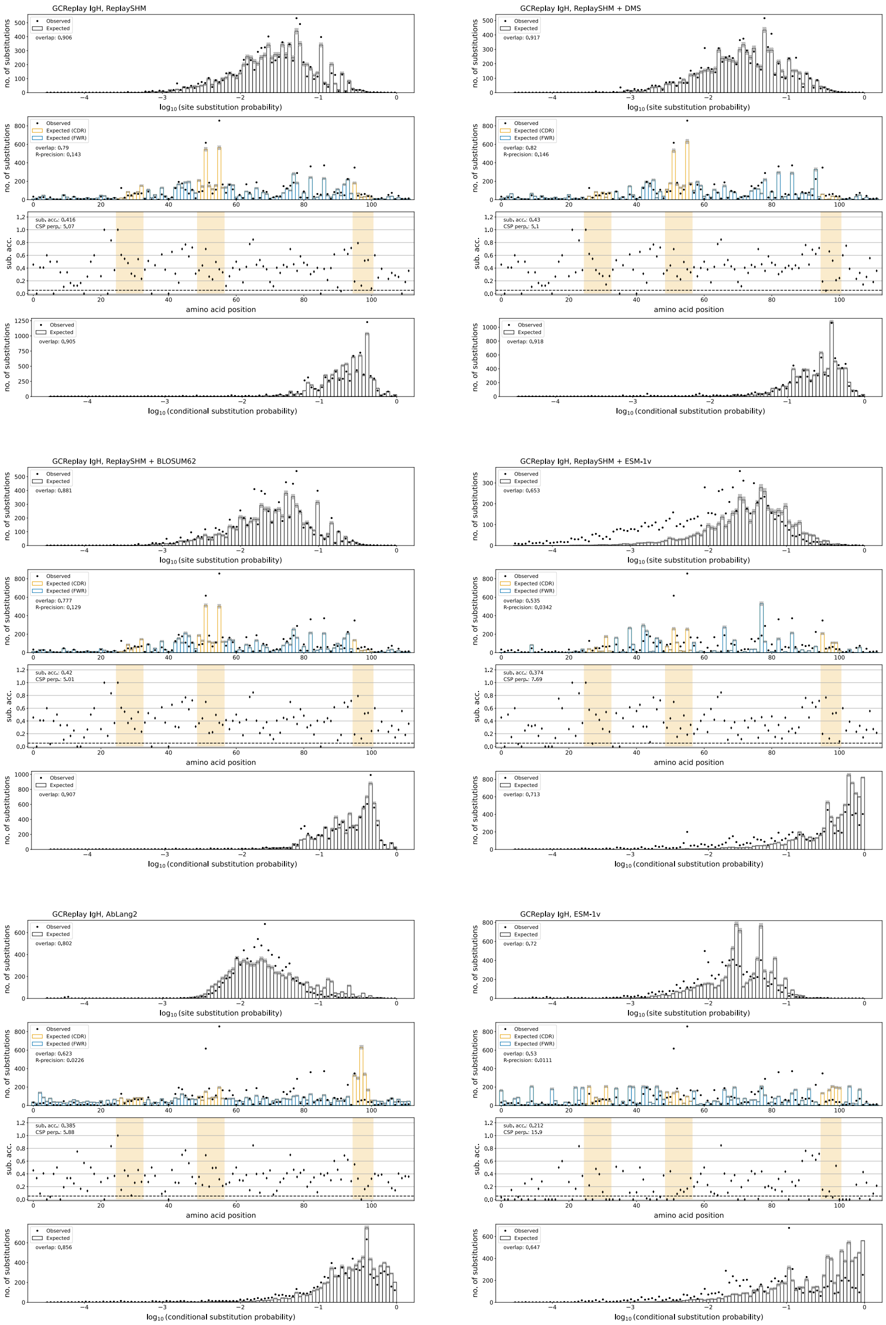


**Fig S8. All models on Replay heavy chain data set.** For each model, there are four panels. The top panel shows the observed and expected number of substitutions where the sites are binned according to their predicted SSP. The second panel shows the observed and expected number of substitutions over site position. The third panel shows the per-site substitution accuracy. The bottom panel shows the observed and expected number of substitutions at sites of substitution, and the sites are binned according to their predicted CSP. The hatch-filled bands are the standard deviations on the expectations.

fig:gcreplay˙igh˙all

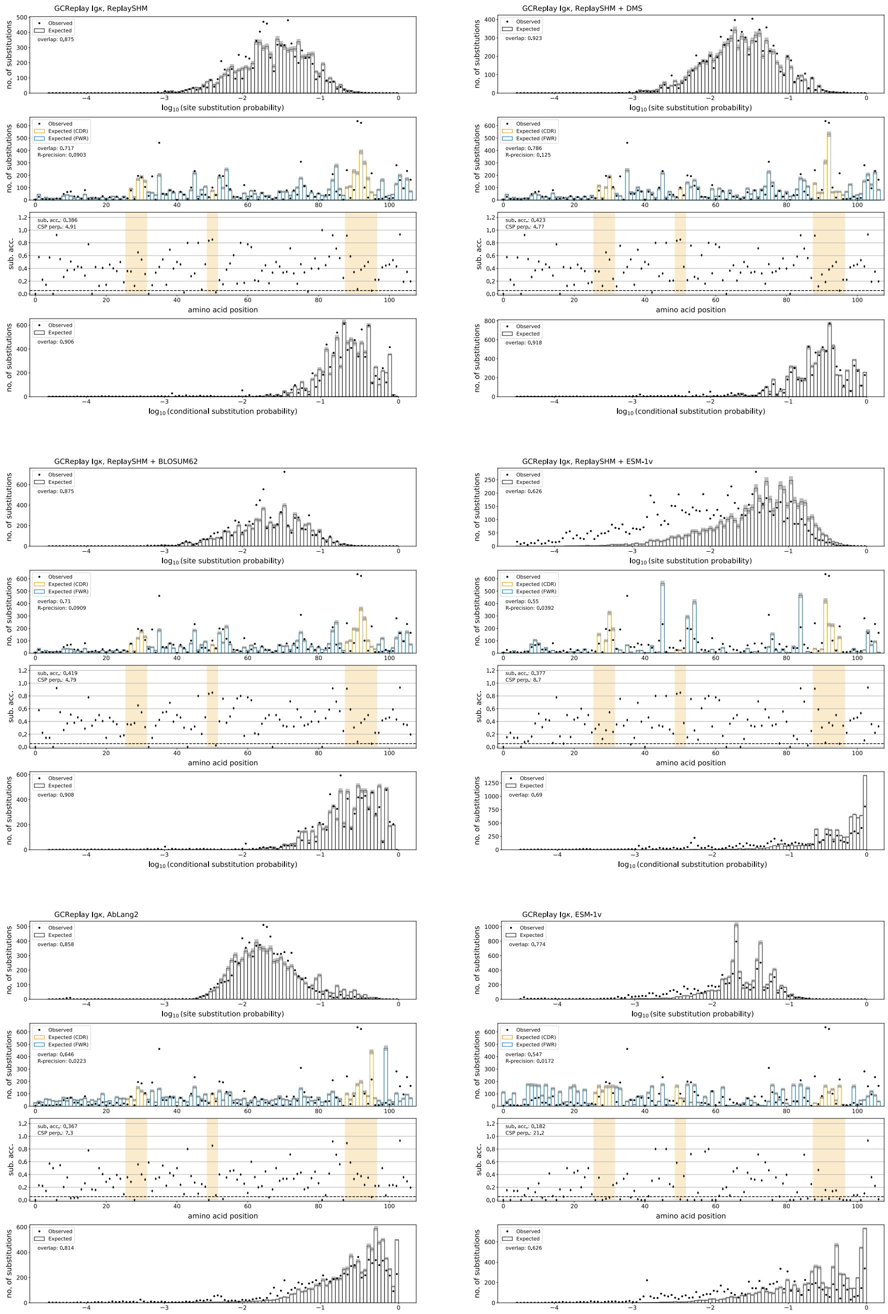


**Fig S9. All models on Replay light chain data set.** For each model, there are four panels. The top panel shows the observed and expected number of substitutions where the sites are binned according to their predicted SSP. The second panel shows the observed and expected number of substitutions over site position. The third panel shows the per-site substitution accuracy. The bottom panel shows the observed and expected number of substitutions at sites of substitution, and the sites are binned according to their predicted CSP. The hatch-filled bands are the standard deviations on the expectations.

fig:gcreplay˙igk˙all

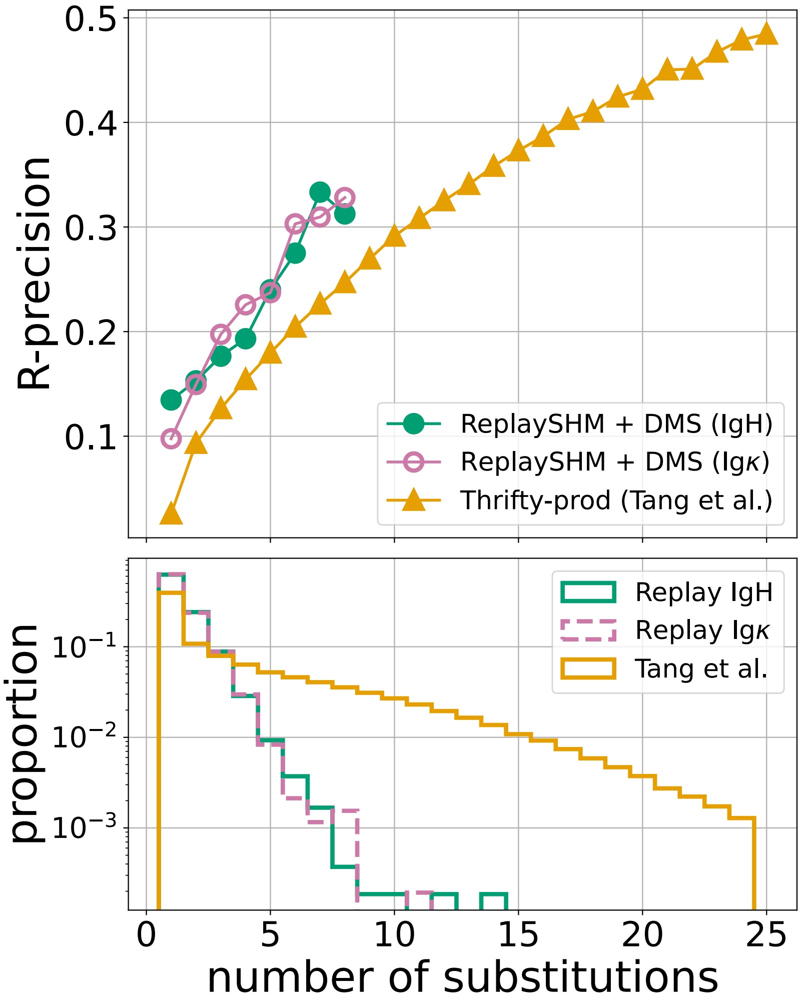


**Fig S10. Comparing results across human repertoires and the Replay data.** (Upper): When stratified by number of substitutions per PCP, R-precision of ReplaySHM + DMS on Replay data is better than models on human repertoire data. The Thrifty-prod model is shown, which is the best model that we evaluated on human data. (Lower): Distribution of the number of substitutions per PCP in the Replay data sets and the Tang et al. data set. The Tang et al. data (and similarly in the other human repertoire data sets) consists of more PCPs with higher number of substitutions than the Replay data.

fig:rprec˙vs˙nmuts

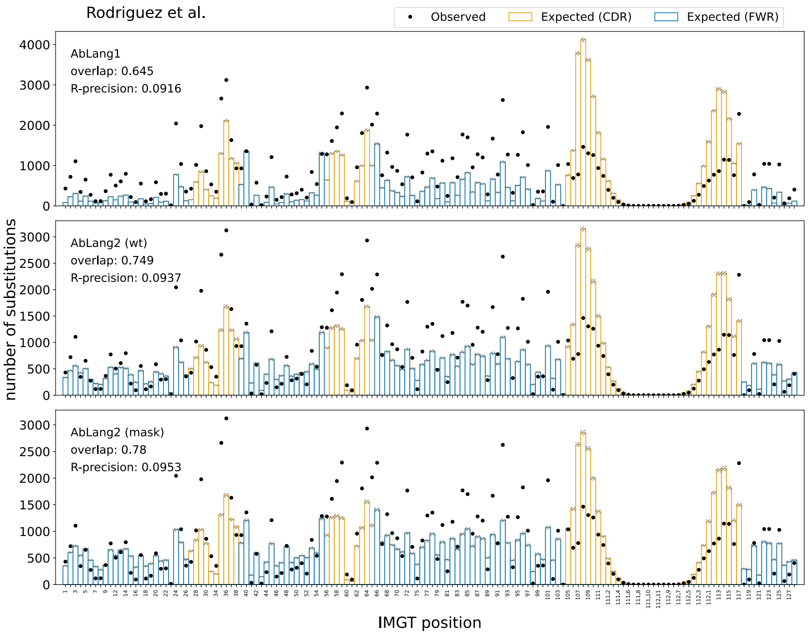


**Fig S11. Improved antibody prediction with AbLang2 on Rodriguez et al. IgH data set.** Results are presented for AbLang1 and AbLang2 using both scoring strategies. AbLang2 (mask) achieved the best performance across all metrics, and was used in all other AbLang2 results presented. fig:ablangs˙comp

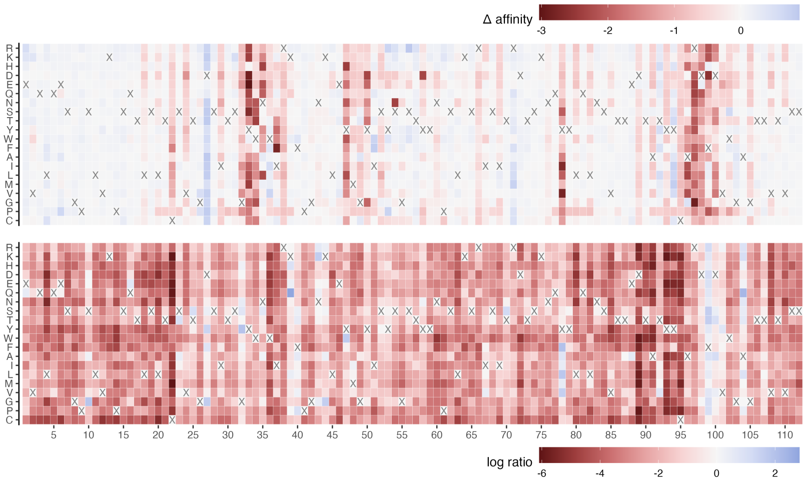


**Fig S12. ESM-1v predictions do not agree with DMS affinity findings in Replay IgH naive sequence.** (Upper): DMS affinity measurements for the Replay IgH naive sequence. (Lower): ESM-1v predictions for the same naive sequence, presented as the log transformed probability ratio. The wildtype amino acid at each site in the naive sequence is depicted with a 'X'. Values are relative to the wildtype amino acid in the naive sequence context.

fig:sel˙heatmaps

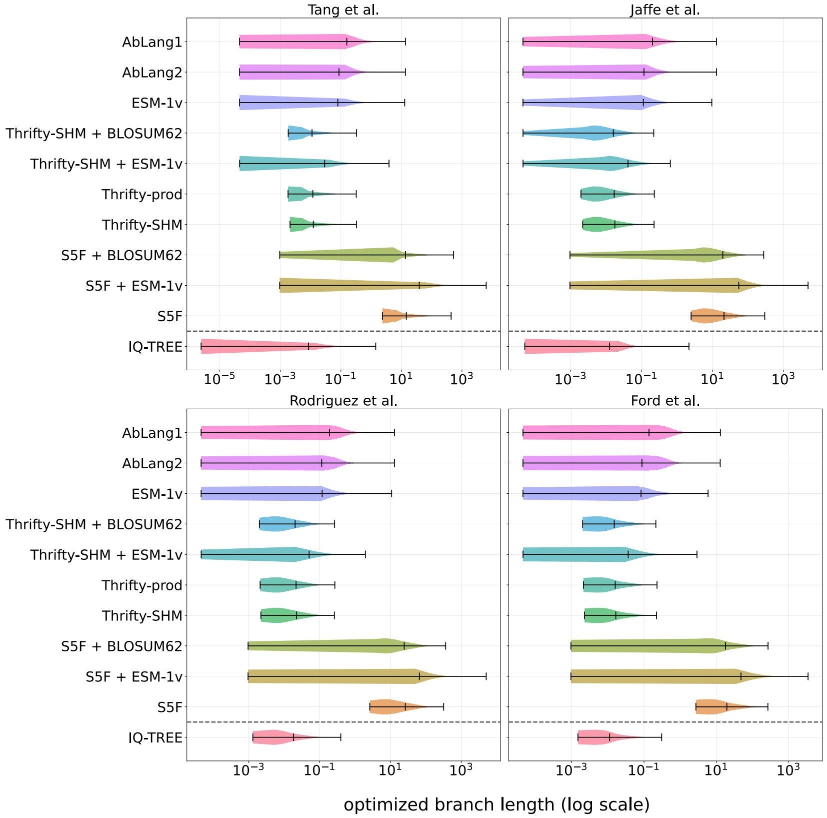


**Fig S13. Optimized per-PCP branch lengths ($\hat{\tau}$) vary widely across models.** The distribution of $\hat{\tau}$ for PCPs under each model is shown for all human data sets. Median $\hat{\tau}$ values are indicated with vertical dashed lines. For reference, the IQ-TREE branch lengths are shown.

fig:opt˙bl

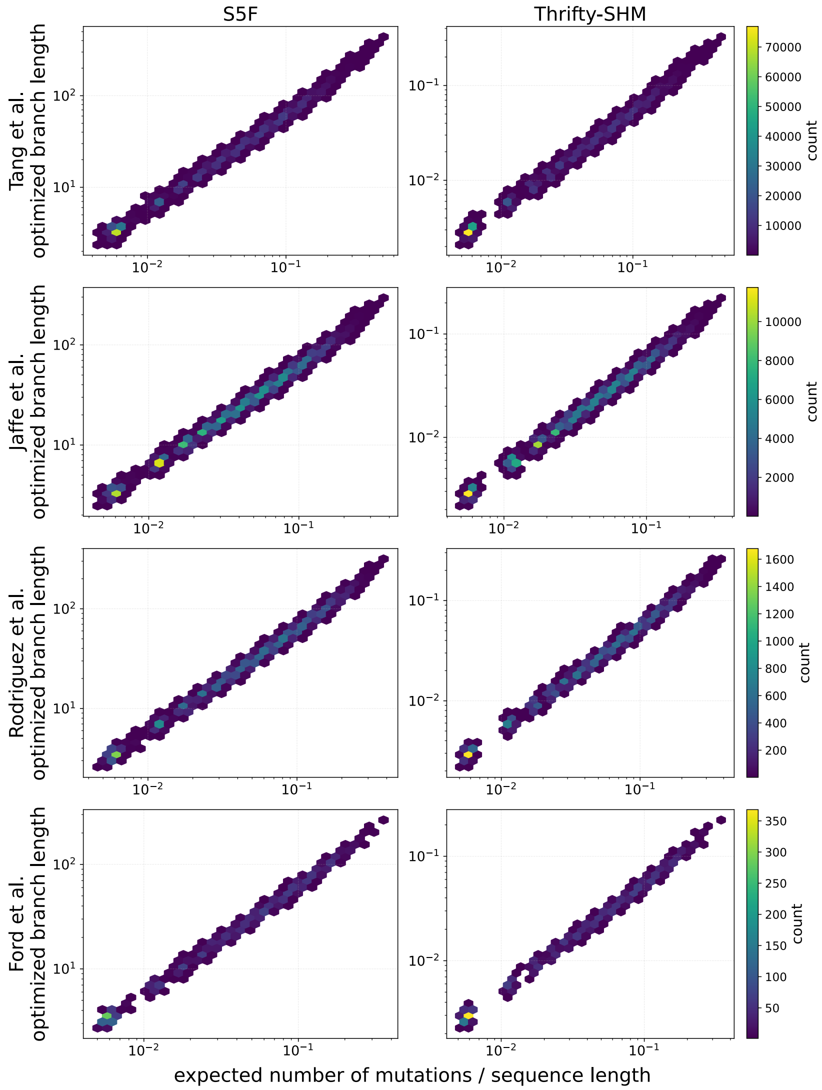


**Fig S14. NT framework models predict mutation to occur at frequencies below one mutation per site.** For all PCPs in the human data sets, the expected number of substitutions per PCP (normalized by codon length) under the S5F and Thrifty-SHM model is plotted against the optimized branch length ($\hat{\tau}$). Colored bins indicate the density of points in the scatter plot.

fig:nt˙mut˙bl

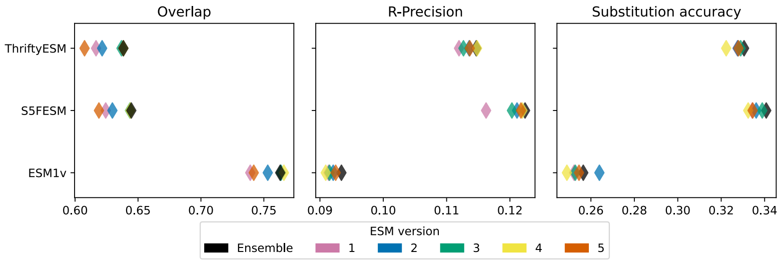


**Fig S15. The ensemble ESM-1v model does not offer a significant performance improvement over individual models in the Rodriguez et al. data set.** For reference, the best-performing model on this data set obtained an overlap of 0.900, R-precision of 0.163, and substitution accuracy of 0.370. This trend was observed in when ESM-1v was used in a standalone model as well as when it was used to compute selection factors in the NT-AA framework. No model consistently performed the best across all metrics. Model 1 of the ESM-1v ensembles was used in all other ESM-1v results presented.

fig:esm˙models
